# DeepReweighting: Reparameterizing Force Field under Explainable Deep Learning Framework

**DOI:** 10.1101/2024.08.07.607110

**Authors:** Junjie Zhu, Zhengxin Li, Junxi Mu, Bozitao Zhong, Haifeng Chern

## Abstract

The analysis and prediction of biomolecule structures is indispensable in various research fields such as protein engineering, mRNA vaccine research and targeted drug screening. Molecular dynamics (MD) simulation is currently the most commonly used and reliable tool for sampling dynamic conformation ensemble of large biomolecules. However, the force fields commonly used in MD simulations are composed of empirical parameters, and their accuracy are limited by the training set data used for parameterizing the force field.In response to these challenges, we have developed a novel force field optimization strategy based on an explainable deep learning framework, DeepReweighting, for rapid and precise force field re-parameterization and optimization. DeepReweighting demonstrates a significant increase in re-parameterization efficiency compared to traditional Monte Carlo method and exhibits greater robustness. Furthermore, DeepReweighting can rapidly re-parameterize any existing or custom differentiable parameters in the force field, providing a faster and more accurate tool for optimizing and utilizing molecular force fields.

## Introduction

The analysis and prediction of biomolecule structures is indispensable in various research fields such as protein engineering, mRNA vaccine research and targeted drug screening. In recent years, numerous computational methods have been developed for predicting crystal structures of biomolecules. AlphaFold2, for example, has demonstrated the capability to predict protein static structures with remarkable precision. Meanwhile, a series of deep learning methods, represented by trRosetta-RNA, have also shown promising performance in predicting RNA structures. However, in functional studies, static structures often prove insufficient. We often require dynamic structural information of large biomolecules to intricately describe the mechanisms through which these molecules exert their functions.

Molecular dynamics (MD) simulation is currently the most commonly used and reliable tool for sampling dynamic conformation ensemble of large biomolecules. It provides a comprehensive supplement to experimental values in analyzing the structural properties of biomolecules. MD simulations are based on first principles, utilizing force fields and solvation models to calculate the potential energy of molecular-solvent systems. The velocity and acceleration at each moment are then solved for each individual atom, enabling the simulation of dynamic properties exhibited by the system over a period of time. The trajectories generated through MD simulations not only provide direct insights for interpretating molecular function or reaction mechanism, but also serve as valuable reference data to complement various data-driven research methods currently under development.

However, the force fields commonly used in MD simulations are composed of empirical parameters, and their accuracy are limited by the training set data used for parameterizing the force field. Traditional structured protein force fields, such as ff03 and ff14SB, show significant discrepancies in simulating local structural features like J-coupling and chemical shifts for intrinsically disordered proteins (IDPs) compared to real NMR observations. Even with force fields specifically re-parameterized for IDPs, such as ff03CMAP, ff14IDPs and ESFF1, the simulation performance on global features like radius of gyration (Rg) remains suboptimal. For RNA, nearly all existing force fields, including OL3 and BSFF1, struggle to achieve ideal simulation accuracy. Therefore, a rapid and effective reparameterization strategy for force fields plays a crucial role in the development and use of force fields.

Previously, Jing Huang et al proposed a force field parameter reweighting method during the optimization of CHARMM36. This method involves Monte Carlo optimization of perturbed force field energies to reweight the conformation ensemble sampled in MD simulations that conform to the Boltzmann distribution. While this reweighting method allows us to obtain a re-parameterized force field that better aligns with experimental observation, the strong randomness in Monte Carlo reweighting leads to uncontrollable perturbations.

Moreover, to enhance the accuracy and transferability of various force fields, specific modifications to force field parameters for different systems have become a common approach in force field optimization and application in recent years. In these studies, Monte Carlo reweighting has imposed substantial speed limitations on generating personalized force field parameters. Meanwhile, the transferability of reweighting methods is relatively low, requiring a substantial foundation of specialized knowledge for application and understanding.

In response to these challenges, we have developed a novel force field optimization strategy based on an explainable deep learning framework, DeepReweighting, for rapid and precise force field re-parameterization and optimization. DeepReweighting demonstrates a significant increase in re-parameterization efficiency compared to traditional Monte Carlo method and exhibits greater robustness. Furthermore, DeepReweighting can rapidly re-parameterize any existing or custom differentiable parameters in the force field, providing a faster and more accurate tool for optimizing and utilizing molecular force fields.

## Methods

### Force field re-parameterization

For a certain molecular system, we can consider that all conformations are described by corresponding high-dimensional vectors *x*. In molecular dynamics simulations, we can envision *x* as encompassing the Cartesian coordinates of all atoms within the conformation, and the potential energy *E*_*λ*_(*x*) of these conformations can be calculated with a particular force field parameterized by *λ*. Ideally, the conformations *x* conform to Boltzmann distribution in thermodynamic equilibrium state at temperature T, as expressed in Equation 1.

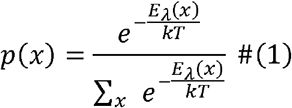

Here *p*(*x*) denotes the probability of conformation *x*, and *k* is Boltzmann constant.

Actually, the force field potential energy *E*_*λ*_(*x*) that we compute serves as an estimate for the true molecular potential energy *E*_0_(*x*). Since in molecular dynamics simulations, we iteratively and non-independently sample configurations based on *E*_*λ*_(*x*), discrepancies between *E*_*λ*_(*x*) and *E*_0_(*x*) can lead to significant differences between simulation results and experimental observations. To minimize such disparities, it is common practice to reparameterize force field parameters based on experimental data or high-precision quantum mechanical calculations in the hope of aligning the ensemble-averaged observables of the simulated system more closely with experimental observations.

Based on the aforementioned motivation, Jing Huang et al. introduced Monte Carlo reweighting for optimizing force field parameters in the development of CHARMM36m. The core idea involves perturbing the CMAP parameters of the force field, redistributing configurations based on energy, and employing Monte Carlo optimization to drive changes in CMAP parameters. This process aims to make the ensemble of redistributed conformations more reasonable in terms of left-handed helical content.

Reparameterization of the force field based on reweighting relies on two major assumptions.

i. The parameters to be optimized must be correlated with target experimental values. For example, correlations have been observed or demonstrated in prior studies between CMAP and the left-handed helical proportion and dihedral angle distribution in proteins, as well as between the ε parameter in solvent models and radius of gyration (Rg).
ii. Reweighting requires a well-sampled ensemble base generated by the original force field that adheres to the Boltzmann distribution. This necessitates a sufficiently large sampling range in simulations, providing a good estimate of the thermodynamic equilibrium state (or the denominator part of Equation 1).

Under the two assumptions, we can precisely describe the relationship between changes in force field parameters and ensemble-averaged observables using the following equation.

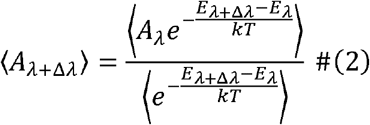

Here ⟨*A*⟩ represents the ensemble-averaged physical quantity of interest during the reparameterization process. *λ* and *λ* + Δ*λ* denote the force field parameters *λ+* Δ *λ* before and after optimization respectively.

### DeepReweighting

In this work, we employ DeepReweighting for stable and rapid force field reparameterization based on reweighting. Utilizing Equation 2, we have developed an optimization algorithm based on matrix calculations, with the force field parameter change (Δ*λ*) treated as a trainable parameter. We update the force field parameters using gradient descent. Taking reparameterization of Lennard-Jones potential in solvent model with respect to Rg as an example, we can transform Equation 2 as follows:

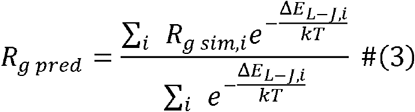

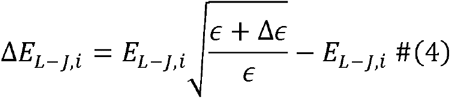

Where *R*_*g pred*_ is the predicted Rg under the new L-J potential parameters, and *R*_*g sim,i*_ is Rg of the i-th conformation in the simulation. Since we are adjusting the □ parameter in the L-J potential, we represent Δ*E* _*L*−*J,i*_ as a function of *Δϵ*. Using Equations 3 and 4, we can treat the optimization problem for the *ϵ* parameter as a process similar to updating a neural network in deep learning. In the training process of a neural network, we iteratively update the network parameters based on their gradient on loss function, aiming to minimize the loss between network prediction and ground truth. In this context, our goal is to adjust the *Δ ϵ* parameter to make *R*_*g pred*_ closer to the experimental value *R*_*g exp*_. Therefore, we can use L1 loss as follows to supervise optimization of *Δϵ*:

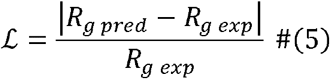

So far, we can define a deep learning framework for optimizing Δ ϵ. Its input contains Rg of each conformation in base trajectory (the focused physical quantity) and the LJ potential energy (used to calculate Δ*E*_*L*−*J,i*_ based on *Δϵ*). The trainable parameters of the network are composed of *Δϵ*, and the training of the network is supervised by the L1 loss from Equation 5. We implement this process using PyTorch and utilize CUDA for GPU acceleration in the optimization process. Tests on various force field parameters and target physical quantities are conducted as detailed in the results section.

## Results

### Reparameterizing solvent model

We tested DeepReweighting on optimizing parameter *ε* in solvent models across 28 IDP systems, comparing it with traditional MC Reweighting. The MD trajectories of these 28 systems were previously validated for convergence in the solvent model development by Junxi Mu et al, indicating sufficient sampling.

We first compared the reweighted ensembles’ radius of gyration (Rg) and the optimized ε parameters from both methods (Figure 1A). While both optimization methods showed comparable Rg predictions across all systems, *ε* obtained through reparameterization were quite different on several systems, indicating that the two methods converged to different local minima during optimization. This also suggests that *ε* and Rg do not necessarily follow a linear relationship as suggested in previous studies. The mean *ε* obtained from DeepReweighting was slightly higher than that through MC Reweighting, while Rg predictions were consistent.

**Figure 1.**
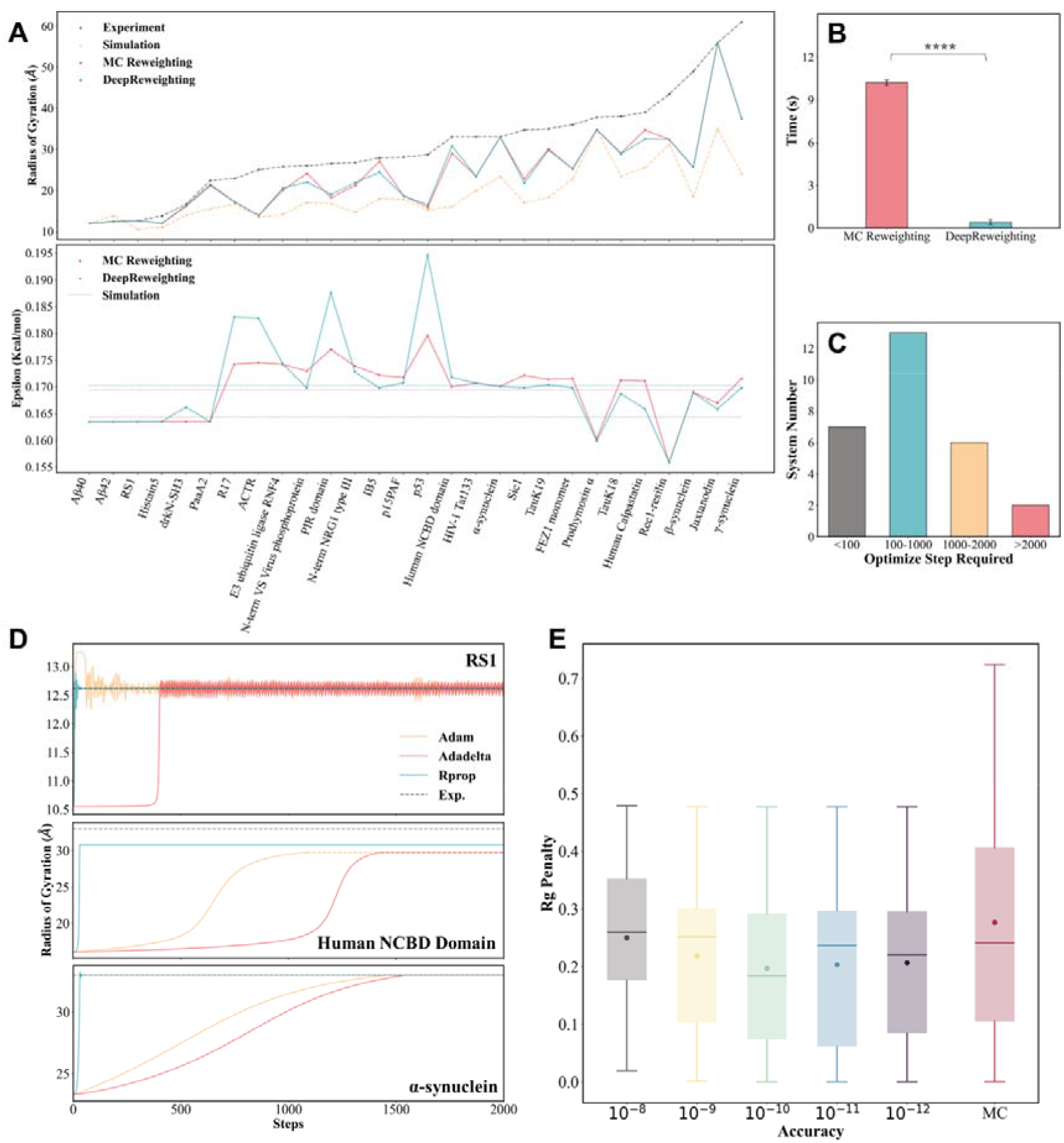
Results on Reparameterizing solvent model. (A) DeepReweighting-predicted Rg and compared to MC Reweighting. (B) Optimization speed of both methods. (C) Convergence of DeepReweighting. (E) Impacts of different model optimizers. (F) Impacts of different numerical precision.

**Figure 1.**
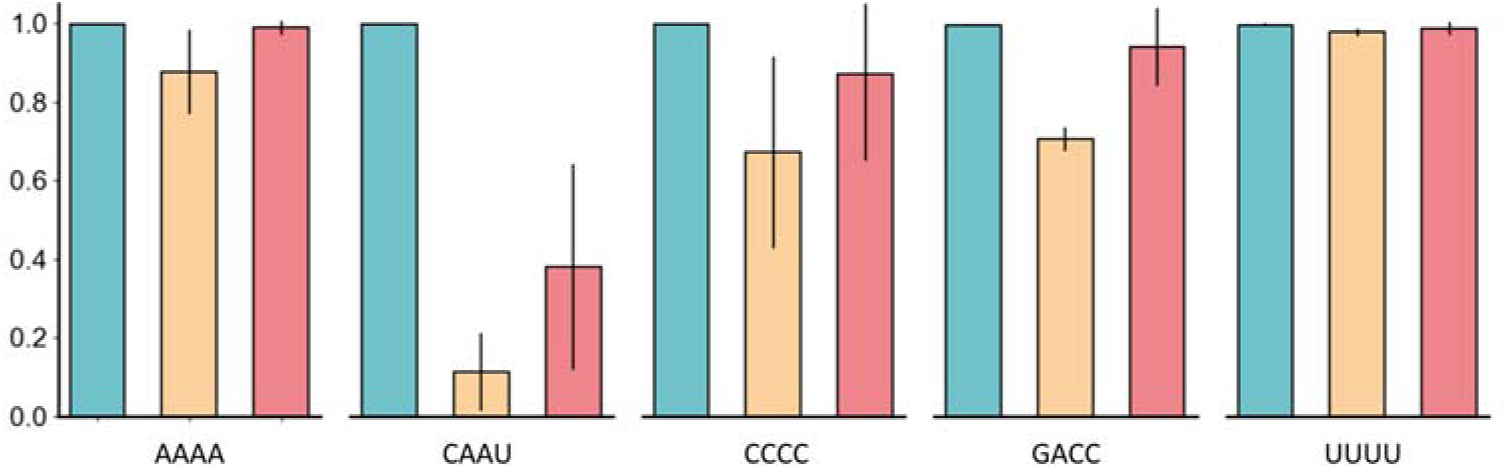
Results of non-intercalated conformation ration of RNA tetranucleotides simulated with DeepReweighting-Reparameterized model(Green), CMAP1(yellow), and OL3(red).

Next, we separately recorded the runtime required for 1000 optimization steps using DeepReweighting and MC Reweighting on Aβ40. The results indicate that DeepReweighting is more than 20 times faster on average than MC Reweighting (Figure 1B). Since we based the optimization on 10,000 frames of Rg and protein-solvent LJ potential for each tested system, the optimization times for both methods are essentially the same across different systems. We conducted three repetitions of tests using seven different optimizers within the DeepReweighting framework and three repetitions for MC Reweighting. The two methods exhibited a significant difference in paired t-tests (*p* ≪ 0.000.1).

Furthermore, we tested the minimum optimization steps required for DeepReweighting to converge across all test systems. We introduced EarlyStopping module from PyTorch to monitor the convergence of the loss function and recorded the stopping optimization steps (Figure 1C). Due to the difficulty in quantitatively assessing the convergence of MC Reweighting, we did not make direct comparisons here. In previous studies MC Reweighting often require 1,000 or 2,000 steps. The results indicate significant variability in the number of optimization steps required for DeepReweighting to achieve convergence across different systems. In systems such as Abeta40 and RS1, where the reweighting results are highly stable, DeepReweighting often converges in just a few dozen steps. In contrast, TauK18 requires approximately 2,400 steps to reach convergence. In general, reweighting for almost all systems converges within 2,000 steps, meaning that DeepReweighting typically completes the reparameterization of a solvent model for a system in less than 1 second on CPU. This operational efficiency is significantly higher than that of MC Reweighting.

We also tested the impact of hyperparameter settings on the model’s performance, including the choice of optimizer and numerical precision for calculations. Among all seven tested optimizers, Rprop performed optimally in minimizing the loss function and converge rapidly, demonstrating the ability to navigate through local minima encountered during optimization. Adam, Adadelta, and AdamW tend to get stuck in local minima in unstable systems, while SGD, ASGD, and RMSprop exhibit significant fluctuations and slow convergence in most systems (Figure 1D, Figure S1). Although Rprop was originally designed for global gradient descent optimization and may not be suitable for most current deep learning methods using mini-batch training, it aligns well with our task and performs the best. Therefore, we choose Rprop for optimizing DeepReweighting. Regarding numerical precision, we recorded the performance of DeepReweighting when using floating-point numbers with different levels of precision. The results indicate that the model performs optimally when calculations are carried out with a maximum precision of 10^−10^ (Figure 1E). Moreover, DeepReweighting outperforms MC Reweighting across all tested precisions.

In summary, DeepReweighting is comparable to MC Reweighting in precision and significantly outperforms it in efficiency. Therefore, DeepReweighting is proved to be a highly suitable tool for reparameterizing solvent models.

### Reparameterizing RNA force field

RNA plays a crucial role in life functions, including the transmission of genetic information through transcription and translation (mRNA), and various physiological functions such as regulation and catalysis (ncRNA). The structure and dynamic conformational ensemble of RNA are essential for its functionality. Current experimental methods for resolving RNA structures are expensive, significantly hindering research and development of RNA structure-function relationships. In this context, molecular dynamics (MD) simulations become an essential research tool, with the accuracy of these simulations largely determined by the molecular force fields used. Existing RNA force fields have several issues, including the propensity to produce intercalated conformations.

Previous studies have aimed to improve RNA force field accuracy, such as the cmap1 force field developed by Chenjun et al., which introduced corrected CMAP parameters using Monte Carlo annealing simulations combined with a reweighting algorithm to reduce the proportion of intercalated conformations. However, their method has significant room for improvement, including the slow speed of Monte Carlo annealing simulations and the perturbation of unrelated regions, which introduces errors.

RNA tetranucleotides, due to their small structure, low computational cost, and inclusion of most RNA intramolecular interactions, as well as abundant experimental data, have become a standard system for testing RNA force fields. In this study, we used a deep reweighting algorithm to adjust CMAP parameters to address the inaccuracies in simulating RNA tetranucleotides with current force fields, which tend to produce intercalated conformations.

To compare with the previous Monte Carlo annealing simulation algorithm, we followed the same workflow as Chenjun et al.: first, we calculated and analyzed the proportion of intercalated conformations and the corresponding zeta-alpha dihedral angle distributions for five tetranucleotides (AAAA, CAAU, CCCC, GACC, UUUU) in the OL3 force field simulation trajectories. We then used the deep reweighting algorithm to develop CMAP parameters for the zeta-alpha dihedral angles, which were subsequently introduced into the force field for simulation. We conducted three parallel trajectories of 2 microseconds each with both the OL3 and DR force fields, maintaining consistent temperature, ion conditions, and water box settings with previous work.

Analysis of the simulation results in Fig2 shows that the DR force field effectively suppressed the proportion of erroneous intercalated conformations, significantly improving the simulation accuracy of the RNA molecular force field. This improvement is especially noticeable in the CAAU system, where both the OL3 and CMAP1 force fields performed poorly. The proportion of intercalated conformations in all systems using the DR force field was below 1%, and even zero in two parallel trajectories of the AAAA system, demonstrating the method’s high efficiency and accuracy.

Moreover, analyzing the CMAP parameters themselves reveals that the DR algorithm is more efficient. In Chenjun et al.’s work, the Monte Carlo annealing simulation algorithm took more than ten hours to optimize the CMAP parameters, while the DR algorithm reduced this time to under 10 seconds, a more than 3,000-fold increase in efficiency. The DR algorithm is also more precise; statistical analysis by Zhengxin Li et al. identified that the main defect in the force field leading to erroneous intercalated conformations is the unreasonable energy surface of the zeta-alpha angles in the regions [-30∼-60, -30∼-60] and [3060, 3060], with erroneous intercalated conformations concentrated in the [30-60, 30-60] region and non-intercalated conformations in the [-30∼-60, -30∼-60] region. Raising the energy surface of the [30-60, 30-60] region and lowering that of the [-30∼-60, -30∼-60] region is key to tuning the force field. Both CMAP1 and DR can capture this, indirectly proving the accuracy of the DR algorithm. Additionally, the DR algorithm minimizes perturbations to other unrelated regions, reducing unnecessary errors.In conclusion, the DR algorithm is an important method for developing RNA molecular force fields and can significantly aid in understanding the dynamic structure-function relationships of RNA.

